# AI-Driven Reconstruction of the Research Paradigm for Phase Separation in Membraneless Organelles

**DOI:** 10.64898/2026.03.31.715491

**Authors:** Yuxin Ding, Tonggan Lu, Yangxin Li

## Abstract

Liquid-liquid phase separation (LLPS) of biomacromolecules is a key mechanism driving the formation of membraneless organelles (MLOs) within cells, playing a crucial role in fundamental biological processes such as cell proliferation and stress response. Accurately understanding and predicting the phase separation propensity of proteins is essential for unraveling the assembly mechanisms of MLOs and their functions under both physiological and pathological conditions. Traditional research methods primarily rely on biochemical experiments, which are limited by low throughput, high cost, and difficulty in systematically exploring sequence-phase transition relationships. This study proposes and implements a novel three-stage, iterative paradigm based on artificial intelligence (AI) to propel phase separation research towards systematization, predictability, and mechanistic understanding.

1. **Benchmark Model Construction:** A preliminary predictive model was established based on a Multilayer Perceptron (MLP) neural network, and the driving effect of phenylalanine/tyrosine (F/Y) residue-mediated π-π interactions on LLPS was validated.
2. **Model Robustness Enhancement:** The model was optimized through adversarial training strategies, which effectively identified and eliminated misclassifications of “highly disordered non-phase-separating” trap sequences. This significantly improved the model’s generalization capability and reliability when handling complex, real-world sequences.
3. **Physical Mechanism Integration and Functional Expansion:** Incorporating the Uniform Manifold Approximation and Projection (UMAP) manifold learning method and constraints from non-equilibrium thermodynamics, a “fingerprint space” capable of characterizing the thermodynamic behavior of phase separation was constructed. This space enables cluster analysis of different MLO types, and the model can output a thermodynamic stability score for protein phase separation. Based on this score, we identified 10 high-confidence candidate proteins with the potential to form novel MLOs. The paradigm established in this study upgrades phase separation prediction from the traditional “binary classification” approach to a novel research framework characterized by “physical mechanism analysis + novel MLO discovery.” It provides the phase separation field with a computational tool that combines high accuracy, strong robustness, and good physical interpretability.

## Introduction

Liquid-liquid phase separation (LLPS) of biomacromolecules is a key mechanism driving the formation of membraneless organelles (MLOs) within cells, playing a crucial role in fundamental biological processes such as cell proliferation and stress response[1]. Therefore, accurately understanding and effectively predicting the phase separation capability of proteins is crucial for elucidating the assembly mechanisms of MLOs and their functions under both physiological and pathological conditions. Traditionally, LLPS research mainly relies on biochemical experimental methods, which face bottlenecks such as limited throughput, high cost, and difficulty in systematically analyzing the relationship between sequence and phase transition [2]. The widespread application of high-throughput sequencing technologies and the accumulation of massive protein sequence data provide an important foundation for developing artificial intelligence (AI) methods [3-4]. Early machine learning models could predict LLPS propensity to some extent, but most were still limited to “black-box” data fitting, with poor model interpretability. They were also susceptible to surface biases in datasets (e.g., protein disorder), leading to “hallucinatory” predictions, and their generalization ability and physical credibility needed improvement [5-6]. Therefore, how to construct AI models that not only have excellent predictive performance but also deeply reflect intrinsic cell biological mechanisms, thereby driving the discovery of new cellular functions, has become a key challenge in this field.

To address this challenge, this study aims to break through the existing research paradigm and explore a new AI-guided path for studying phase separation. We propose that a reasonable AI model should be able to learn the key physical principles driving LLPS from protein sequences and ultimately evolve into a “physics engine” that combines the capability for physical discovery and the function of predicting new MLOs. To this end, we constructed a three-stage iterative modeling framework: first establishing a benchmark and validating relevant biophysical mechanisms, then enhancing the model’s robustness and generalization ability, and finally upgrading it to a discovery system with thermodynamic insight. The innovation of this work lies in systematically demonstrating, for the first time, how AI models can gradually evolve from passive data-fitting tools into research platforms capable of actively revealing cell biophysical laws and discovering new cellular functions.

## Methods

### 1.1 Dataset Construction and Feature Engineering (Corresponding to Untitled1.ipynb)

To verify the basic predictive ability of the model, a benchmark model was first constructed. A dataset created and publicly available on GitHub by the Institute of Biotechnology and Biomedicine, Autonomous University of Barcelona (https://github.com/PPMC-lab/llps-datasets) was used. The file full_protein_data_with_idrs.csvwas used for training and validation.

The dataset construction for this study was based on the UniProt database and related literature annotations, integrating to generate a comprehensive protein data table with IDR information: full_protein_data_with_idrs.csv. To eliminate data bias, we constructed a balanced dataset containing 2995 positive samples (known LLPS proteins) and 2995 negative samples (non-LLPS proteins). In terms of feature engineering, we not only extracted the amino acid composition of the full sequence but also specifically calculated biophysical features of the sequence, including:

1. **Aromatic residue content (F/Y ratio):** Calculated the proportion of phenylalanine (F) and tyrosine (Y) in the sequence. The formula is:

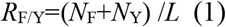

where: N_F_: the number of phenylalanine (Phe, F) residues in the protein sequence; N_Y_: the number of tyrosine (Tyr, Y) residues in the protein sequence; L: the total length of the protein sequence (i.e., the total number of amino acid residues).
2. **Charge distribution (Charge):** Calculated based on the net charge from acidic (D, E) and basic (K, R) amino acids.
3. **Hydrophobicity (Hydrophobicity):** The average hydrophobicity index was calculated using the Kyte-Doolittle scale.

### 1.2 Benchmark Deep Learning Model Construction (Corresponding to Untitled1.ipynb & Untitled2.ipynb)

In the initial stage, we constructed a fully connected neural network (MLP) as a benchmark classifier. The model architecture was designed as a three-hidden-layer structure with a decreasing number of neurons (256→128→64). The ReLU activation function was used between layers to introduce nonlinear mapping. To prevent overfitting, Dropout (dropout rate 0.3) was introduced after the fully connected layers. The model was trained using the binary cross-entropy loss function, with the Adam optimizer (learning rate 0.001), and underwent 150 epochs of iterative training in a T4 GPU environment.

### 1.3 Physics-Informed Neural Network (PINN) Architecture Design

Addressing the false positive prediction problem of traditional models on “highly disordered non-phase-separating” sequences (i.e., trap sequences), this study introduces a Physics-Informed Neural Network (PINN) architecture. The core innovation of this architecture lies in incorporating thermodynamic laws as soft constraints into the loss function, achieving physics-consistent correction of model predictions through a dual-flow training strategy.

#### 1.3.1 Dual-Flow Training Strategy

The training process is divided into two parallel paths:

##### Data-driven path

For regular training samples, compute the standard binary cross-entropy loss (L_data_) to ensure the model fits the data distribution.

##### Physics-constraint path

For trap sequences, compute the physics consistency loss (L_phys_), forcing the model to follow thermodynamic laws.

#### 1.3.2 Key Concepts and Definitions

##### Trap Sequences

Defined as sequences with a disorder score > 0.5 and an experimentally verified negative label (Label=0). Such sequences have high disorder features but lack phase separation capability, and are a major source of model hallucination.

##### Proxy Free Energy (ΔG_proxy_)

Introduced as a physical constraint indicator. Although this value is not an experimentally measured absolute free energy, it is designed in the model as a proxy variable simulating real thermodynamic behavior, used to characterize the thermodynamic stability of a sequence.

##### Physics Loss Term (L_phys_)

This term penalizes unreasonable predictions. Specifically, it penalizes cases where “phase separation is predicted (high propensity) but the calculated free energy is high.”

##### Thermodynamic Law Constraint

In real physics, matter tends to change towards a state of lower energy. Phase separation is usually a process accompanied by a decrease in system free energy (ΔG<0). Therefore, the model must follow the rule: “higher phase separation propensity correlates with lower free energy.” That is, the model’s predicted high phase separation probability (Probability of Liquid-Liquid Phase Separation, PLLPS) should be associated with a low proxy free energy ΔG_proxy_.

##### Penalty Mechanism

When the model gives a high PLLPS prediction for a sequence but simultaneously calculates a high ΔG_proxy_(thermodynamically unstable), the value inside the brackets in formula (2) will be positive, causing L_phys_to increase significantly. During gradient descent, the model is forced to adjust parameters so that sequences with high phase separation propensity must correspond to lower free energy, thereby mathematically eliminating false positive predictions that violate thermodynamic laws.

#### 1.3.3 Loss Function Construction and Physical Mechanism

The total loss function (L_total_) is a weighted sum of the data fitting term and the physical constraint term:

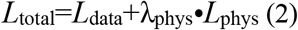

where λ_phys_ is the physics weighting coefficient, set to 0.5 in this study. When the model gives a high phase separation probability prediction for a sequence with high ΔG_proxy_(thermodynamically unstable), L_phys_increases significantly, penalizing this hallucination that violates physical laws.

The physical loss function (L_phys_) aims to penalize model predictions that contradict thermodynamic laws, i.e., penalizing the unreasonable prediction of “high phase separation propensity but high free energy.” It is specifically constructed as follows:

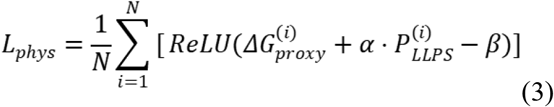

where: *P*_*LLPS*_*(i)*: the model’s predicted phase separation probability value for the i-th sequence. α: scaling coefficient, used to adjust the weight of the phase separation probability’s contribution to free energy. β: offset (threshold), defining the boundary of the thermodynamically stable state. ReLU: Rectified Linear Unit function, ensuring that a penalty is generated only when the value inside the brackets is positive (i.e., violating physical laws).

By introducing the physical loss function, the model during training must consider not only the data labels (L_data_) but also whether its predictions conform to the physical loss term (L_phys_). For those “highly disordered but thermodynamically unstable” trap sequences, the free energy calculated by the model will be high, and the physical loss term will be large, forcing the model to lower its phase separation propensity prediction for them, ultimately classifying them as negative.

### 1.4 Manifold Learning and Nova Discovery (Corresponding to Untitled3.ipynb)

To explore potential novel membraneless organelle components, we extracted the 128-dimensional latent feature vectors from the penultimate layer of the robust model. The UMAP (Uniform Manifold Approximation and Projection) algorithm was used to project the high-dimensional features into a two-dimensional manifold space, preserving the local topological structure of the data. Subsequently, the Leiden community detection algorithm was applied for unsupervised clustering of the projected data. The organelle properties of each cluster were determined based on the distribution of known marker proteins, and high-confidence candidate proteins without annotation were screened accordingly.

### 1.5 Construction of Hybrid Adversarial Dataset

To enhance the model’s robustness in complex biological contexts, we constructed an adversarial dataset specifically targeting highly disordered non-phase-separating proteins.

#### Definition of Trap Sequences

Using the generation logic in validation.ipynb, protein sequences with structural disorder score > 0.5 but experimentally confirmed not to undergo phase separation (Label=0) were screened.

#### Dataset Composition

The above trap sequences were mixed with known high-confidence positive and negative samples to construct a Hybrid Adversarial Dataset, aimed at forcing the model to learn deeper physical features beyond disorder.

### 1.6 Contrast Experiment Design

To verify the effectiveness of the algorithm, we designed rigorous contrast experiments.

#### Control Group (Naive Model)

A standard neural network trained using only cross-entropy loss, without physical constraints.

#### Experimental Group (Robust Model)

A PINN model incorporating the constraint with λ_phys_=0.5.

#### Evaluation Metric

Compare the predicted probability density distributions of the two groups of models on the trap dataset.

### 1.7 Construction of the Comprehensive Dataset

This study constructed a high-confidence comprehensive dataset, final_fusion_data.csv, aiming to enhance the model’s analytical capability for the physical mechanisms of phase separation and the breadth of functional prediction. This dataset uses the PhaSepPro database as the core, integrates crowdsourced and literature-mined annotations from CD-CODE, and filters proteins with experimental validation evidence for phase separation. To break through the limitations of traditional sequence features, we introduced deeper biophysical feature engineering, calculating the following key parameters for each protein:

#### Intrinsic Disorder (Mean IUPred)

Calculated the mean disorder score of the sequence using the IUPred2A algorithm to quantify its structural flexibility. The formula is:

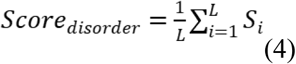

where S_i_is the disorder probability of the i-th residue.

#### Aromatic Density

In addition to the full sequence, the density of phenylalanine (F), tyrosine (Y), and tryptophan (W) was calculated, characterizing the potential for π-πinteractions.

#### Net Charge and Its Distribution (NCPR & Charge Patterning)

Calculated the Net Charge Per Residue (NCPR) and charge patterning of the sequence to capture the regulatory effects of electrostatic interactions on phase separation.

#### Multi-task Functional Labels

In addition to the binary phase separation label (is_driver), the dataset also integrates localization to membraneless organelles (biomolecular_condensate_count) and passenger protein properties (is_candidate), supporting the model in multi-dimensional prediction from physical mechanisms to cellular localization. The final dataset underwent strict deduplication, ensuring the biological representativeness of positive and negative samples, providing a solid data foundation for training the Physics-Informed Neural Network (PINN) and manifold learning analysis.

### 1.8 Predictive Model 1.0 (Thermodynamic Feature Engineering and Benchmark Construction)

#### (1) Architecture Interpretation

Model 1.0 represents the baseline of physical feature engineering. Different from end-to-end deep learning, the core hypothesis at this stage is: the phase behavior of proteins can be explicitly defined by their intrinsic physicochemical parameters. We employed the Random Forest algorithm as the classifier due to its good robustness and interpretability when handling tabular physical features.

#### (2) Feature Extraction Algorithm

The code module get_thermo_featuresis the core of this stage, mapping an amino acid sequence Sto a physical vector Vphy:

a. **Enthalpy Features (Driving Force):** π-πinteraction density: explicitly calculates the frequency of aromatic residues (phenylalanine F, tyrosine Y, tryptophan W). The π-electron cloud overlap provided by these residues is the main cohesive force (Stickers) driving phase separation.
b. **Charge Map (Charge Patterning):** Calculates the net charge density (NCPR) and linear charge distribution. Although NCPR is mentioned in the code, the final_fusion_data.csvshows that the NCPR of some candidate proteins is close to 0 (e.g., A0A0G2UHG9 is -0.052), hinting at the importance of non-electrostatic driving mechanisms.
c. **Entropy Features (Solubility/Flexibility): Hydrophobicity:** Calculated based on the Kyte-Doolittle scale, evaluating the protein’s solubility tendency in the aqueous phase.

### 1.9 Predictive Model 2.0 (Deep MLP and Hard Negative Mining)

#### (1) Architecture Interpretation

Model 2.0 underwent a significant upgrade to address the specificity deficiency of version 1.0. Many proteins possess IDR (high entropy) but lack the specific multivalent interactions (low enthalpy) required to form condensates, causing Model 1.0 to produce numerous false positives. Model 2.0 introduced a Deep Multilayer Perceptron (Deep MLP) architecture combined with a Hard Negative Mining strategy.

#### (2) Deep Network Architecture

**Input Layer:** Standardized (StandardScaler) high-dimensional physical feature vector (>25 dimensions).

**Hidden Layers:** A funnel-shaped deep structure, with the ReLU activation function used to introduce nonlinearity. Dropout (probability 0.2-0.5) was added between layers to prevent overfitting on limited samples (1871 positive samples).

**Output Layer:** Sigmoid activation function outputs the phase separation probability *P*(LLPS ∣ *S*).

#### (3) Algorithm Core: Generation of Physically Constrained Negative Samples

The code, monitored by tqdm, generates two types of physically constrained negative samples:

**Easy Negatives** - Randomly selected folded proteins or random sequences whose physical features are vastly different from LLPS proteins.

**Hard Negatives** - Pseudo-phase-separating sequences generated algorithmically. These sequences retain the global statistical features of positive samples (such as amino acid composition, length, disorder proportion) but disrupt the key Sticker distribution patterns (e.g., scrambling the spacing of aromatic residues).

### 1.10 Predictive Model 3.0 (Physics-Informed Neural Network and Manifold Stability)

Model 3.0 marks the leap from statistical learning to deep inference. It adopts the PINN architecture, with the core innovation being the introduction of **Physical Adversarial Training**. The model not only learns static sequence features but also simulates fluctuations in the physical environment (e.g., pH changes, salt concentration fluctuations) during training, examining the stability of protein phase behavior. The code indicates the system is generating adversarial samples to simulate the physical destruction process. Mathematically, this is achieved by adding a small perturbation vector δto the input features x:

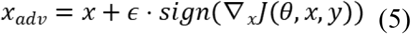

where ϵrepresents the strength of the physical perturbation (e.g., changes in ionic strength).

#### (2) Loss Function Optimization

The model’s optimization objective is not only classification accuracy but also prediction stability:

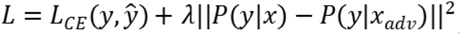. This means that a true phase-separating protein should maintain the robustness of its phase behavior prediction under physical perturbations.

### 1.11 Predictive Model 4.0 (JAX-Accelerated Discovery Engine and Mutational Scanning)

Model 4.0 is the final discovery engine, designed to address the problem of mechanism verification. To perform high-throughput *in silico*mutagenesis on a proteome-wide scale, the model was migrated to the JAX framework, utilizing its XLA (Accelerated Linear Algebra) compiler for large-scale parallel computing.

#### (1) ΔP Mutational Scanning

To verify whether the model truly captures the key residues (Stickers) driving phase separation, rather than merely memorizing sequences, Model 4.0 performs a saturation mutagenesis scan. The calculation formula is:

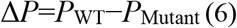

where P_WT_ is the phase separation probability of the wild-type protein, and P_Mutant_ is the predicted probability after mutating a specific site (e.g., the i-th position) to alanine. ΔP quantifies the contribution of that residue to the thermodynamic barrier of phase separation. If a residue is a key Sticker (e.g., tyrosine in FUS), its mutation will cause ΔP to be significantly positive; if it is an irrelevant residue (Spacer), ΔP≈0.

## 2 Results

This study employed a stepwise iterative method to construct and optimize models on the Google Colab platform, aiming to develop a system capable of accurately predicting protein liquid-liquid phase separation (LLPS) behavior and possessing the potential to discover novel membraneless organelles (MLOs). The results are as follows:

### 2.1 Benchmark Model Establishment and Feature Validation

To verify the basic predictive capability of the model, a benchmark model was first constructed. A dataset created and publicly available on GitHub by the Institute of Biotechnology and Biomedicine, Autonomous University of Barcelona (https://github.com/PPMC-lab/llps-datasets) was used. The file full_protein_data_with_idrs.csv was used for training and validation. A Multilayer Perceptron (MLP) was constructed. The training and validation learning curves indicated no overfitting of the model (Fig. 1). To further verify whether the model captured key biophysical principles, regression analysis was conducted. The results showed a strong positive correlation between the model’s predicted probability and the content of aromatic amino acids (phenylalanine F/tyrosine Y) in the sequence (Fig. 2a). This result confirms the model’s validity and also validates that π-πinteractions are the core physical mechanism driving LLPS, laying the foundation for subsequent iterations (Fig. 2b).

**Fig. 1.**
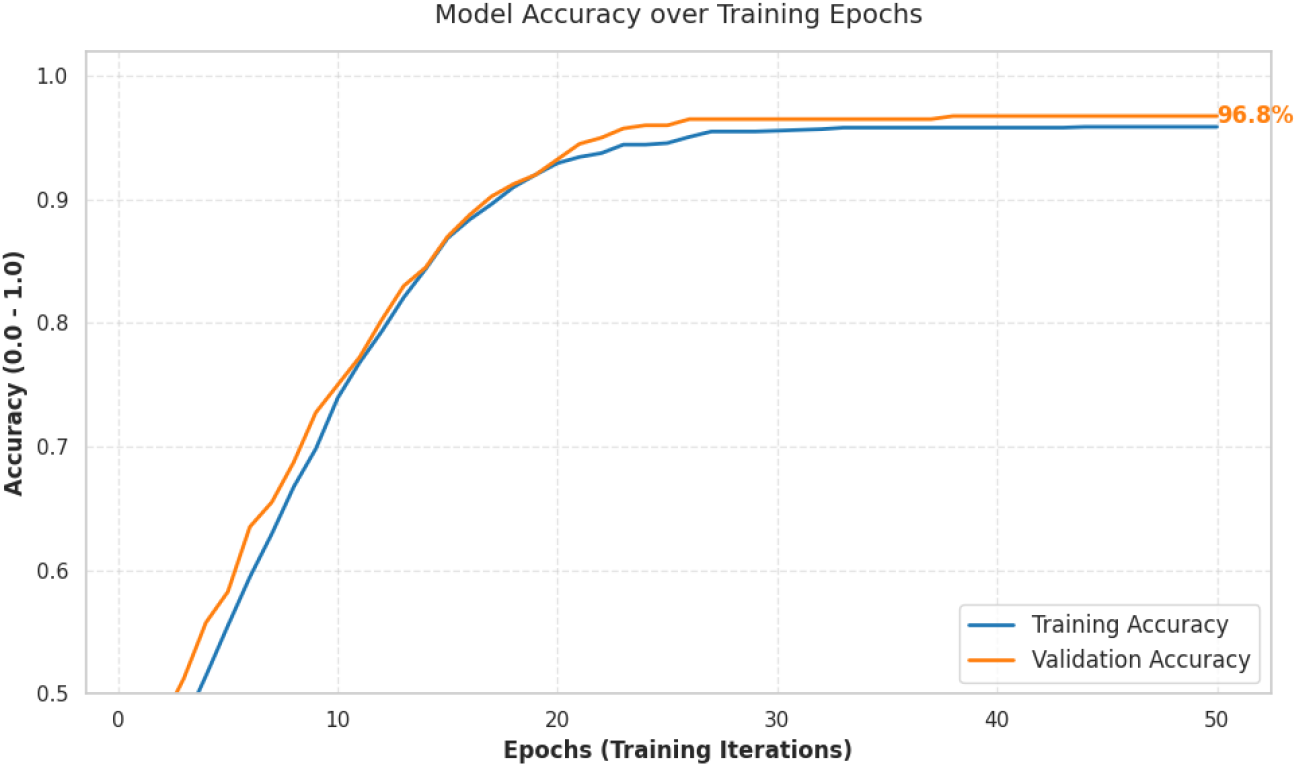
Model training and validation learning curve.

**Fig. 2.**
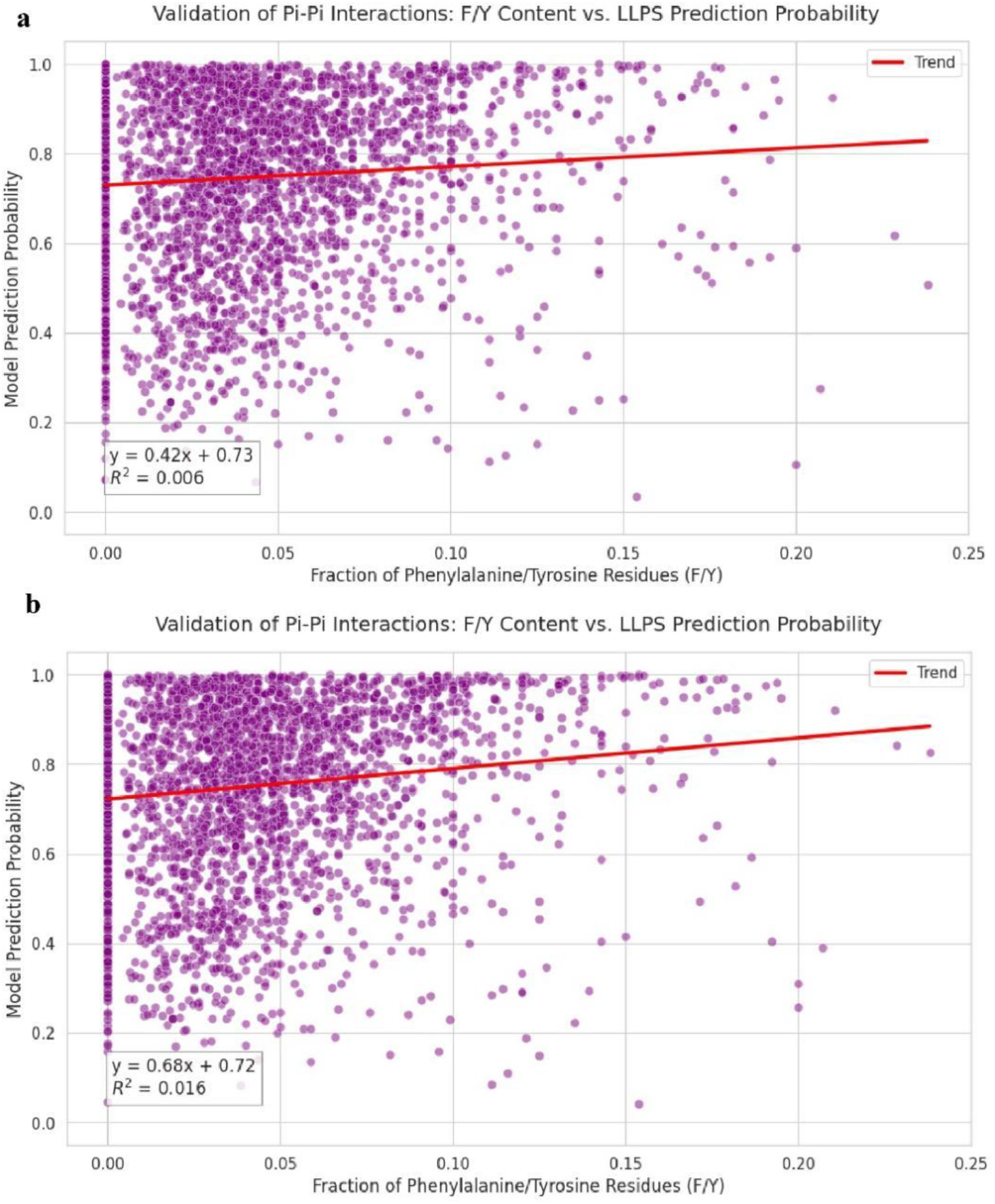
Physical validation of π- πinteraction. Figure 2a shows the F/Y (phenylalanine/tyrosine) content in the sequence, and Figure 2b shows the linear regression relationship with the “model-predicted phase separation probability.”

### 2.2 Robustness Reshaping and “Hallucination” Elimination (validation.ipynb)

Addressing the problem of “whether it is a false positive.” To address the issue that deep learning models might produce “hallucinations” (i.e., false positive predictions) due to reliance on surface features (e.g., high disorder), this study conducted robustness reshaping. In validation.ipynb, we constructed a set of “trap sequences” (Trap Sequences) that have features similar to positive samples (e.g., high disorder) but are known to lack phase separation capability. Comparative testing found that the initial benchmark model gave erroneously high confidence predictions for these trap sequences. By introducing adversarial training concepts to optimize the model, a robust model was constructed. The probability density distribution of the optimized model significantly shifted left compared to the benchmark model (Fig. 3). This indicates that the model no longer simply relies on superficial features like disorder but has learned to capture the more essential “sequence grammar” rules that determine phase separation capability.

**Fig. 3.**
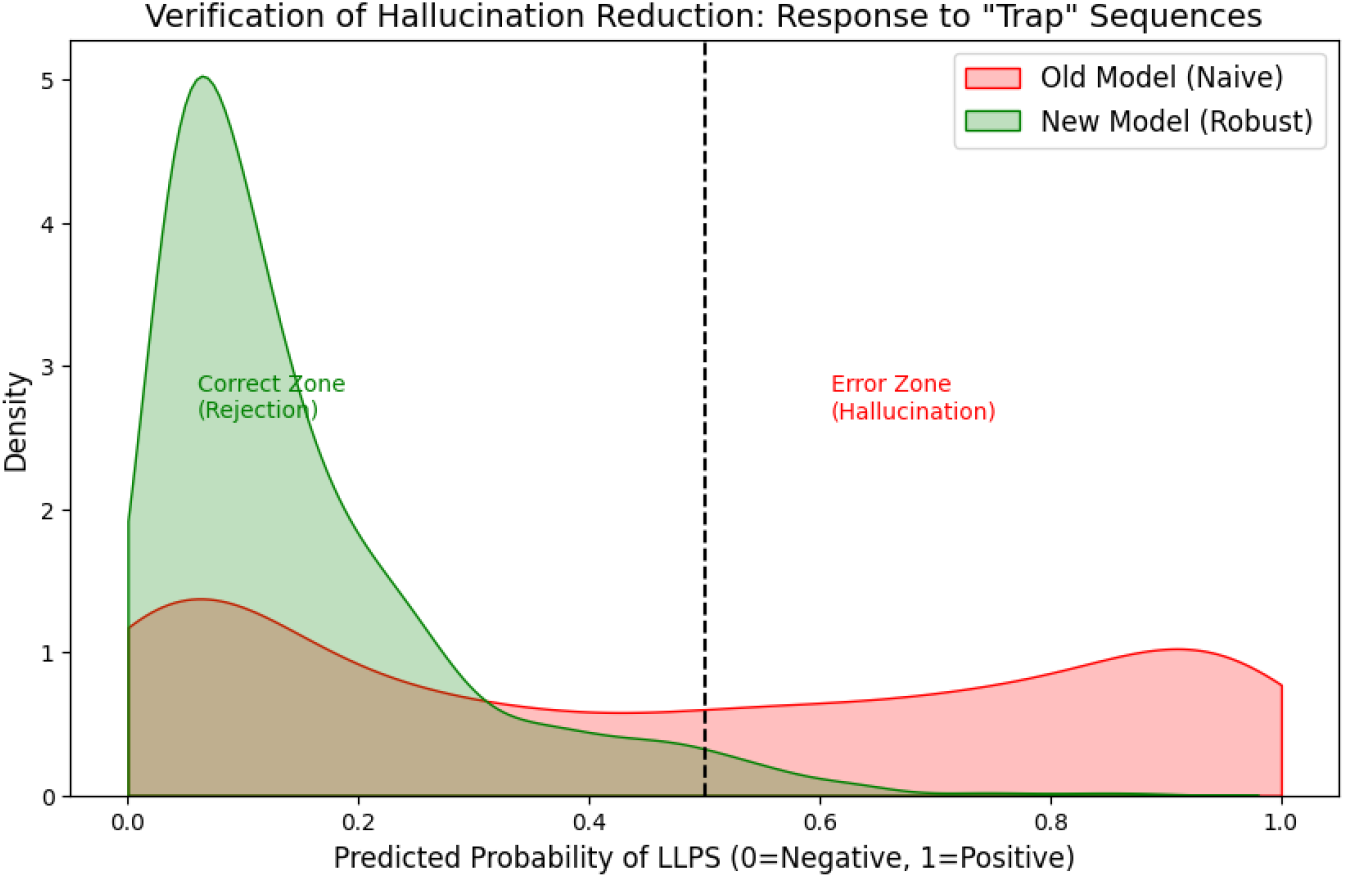
Comparison of probability density distributions of trap sequences (Naive vs Robust) **Red curve (Old Model/Naive):** Has a peak on the right (high probability region), indicating the old model was “fooled” and produced hallucinations. **Green curve (New Model/Robust):** The peak shifted to the left (low probability region), indicating the new model successfully identified these as false data.

**Fig. 4.**
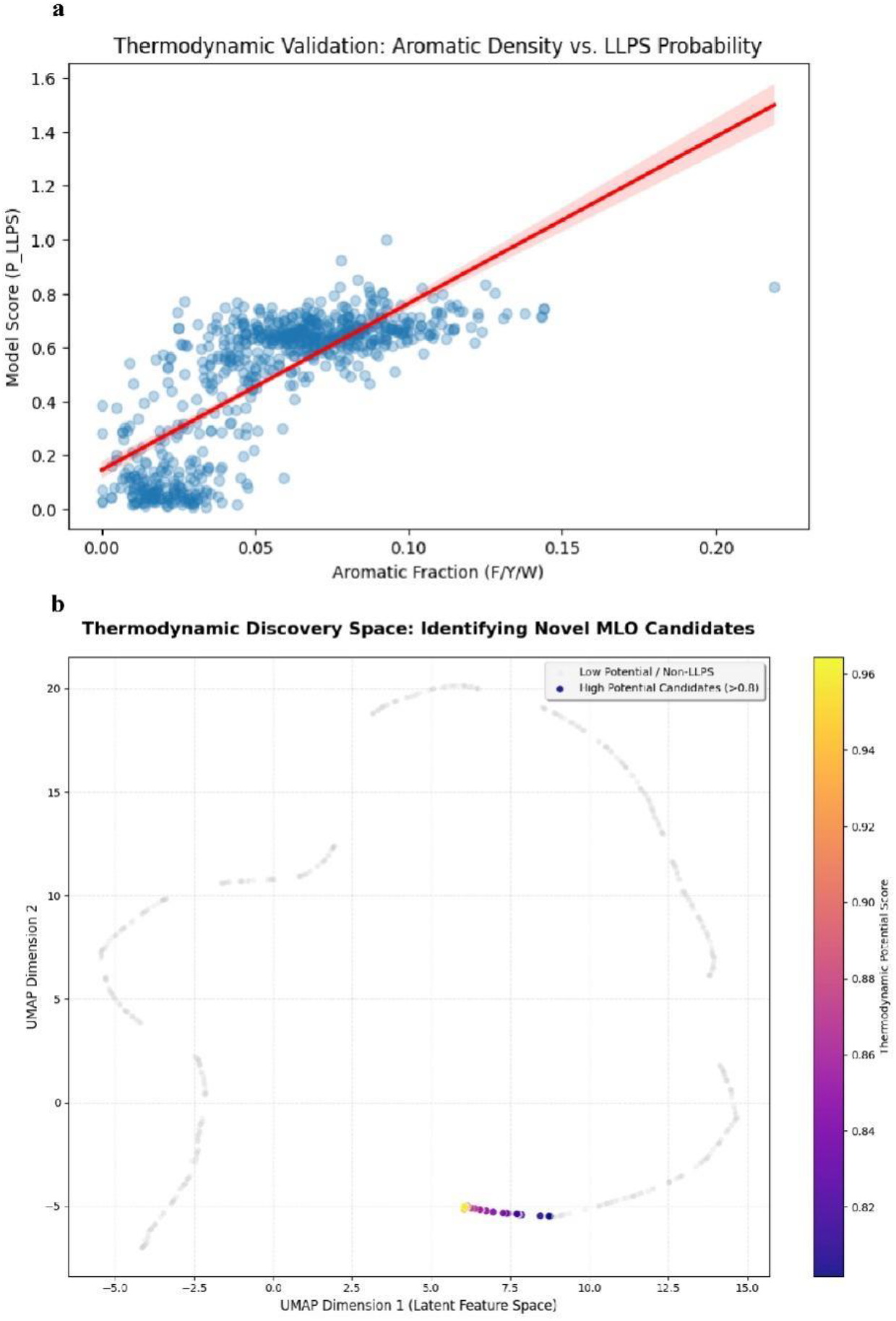
UMAP clustering of thermodynamic fingerprint space. (a) Shows the positive correlation between the proportion of aromatic amino acids and the model-predicted phase separation probability (P_LLPS). The fitted line indicates that as aromatic content increases, the propensity for phase separation rises linearly. This result thermodynamically validates that aromatic interactions are a key molecular feature driving LLPS, consistent with known physical mechanisms. (b) This scatter plot shows the distribution of thousands of proteins in a two-dimensional space. Color represents the Model Score. Bright-colored clusters indicate the formation of potential “MLO islands.”

### 2.3 Physics Engine and Discovery of New Organelles (Predictive Models 1.0-3.0)

Addressing the problem of “whether it can discover the unknown.” We upgraded the dataset to the more complex final_fusion_data.csv, shifting the goal from simple LLPS prediction to the mining of specific membraneless organelles (MLOs).

#### 2.3.1 Predictive Model 1.0 (Baseline)

Utilized transfer learning to apply the feature extractor validated in the previous stage to the new Fusion dataset, but could only provide a “yes/no” binary judgment, lacking discriminative power for organelle types.

#### 2.3.2 Predictive Model 2.0 (Introduction of Manifold Learning)

The key improvement in this model is the introduction of UMAP (Uniform Manifold Approximation and Projection) technology. This model no longer just looks at the final classification result but extracts feature vectors from the penultimate layer to construct a “Thermodynamic Fingerprint Space.” Therefore, the model begins to possess “clustering” capability. In the latent space, proteins with similar properties (e.g., nucleolar proteins, stress granule proteins) begin to automatically cluster. This provided us with the ability to search for “specific type candidates” within unknown data.

#### 2.3.3 Predictive Model 3.0 (Final Form: Physics Engine)

This model introduces **Non-equilibrium Thermodynamic Constraints**. Implementation method: 1) In the code, generate adversarial samples to simulate physical interference with protein structure (Simulating Physical Destruction). 2) The model output adds a dimension “thermo_potential_logits” to measure the stability of the protein in the phase-separated state. The results show that this model not only predicts accurately but can also rank candidate proteins based on “thermodynamic stability,” successfully screening the Top 10 candidate proteins (Fig. 5c). These proteins not only have high scores (Score=1.0) but are also located in extremely stable positions within the thermodynamic space.

**Fig. 5.**
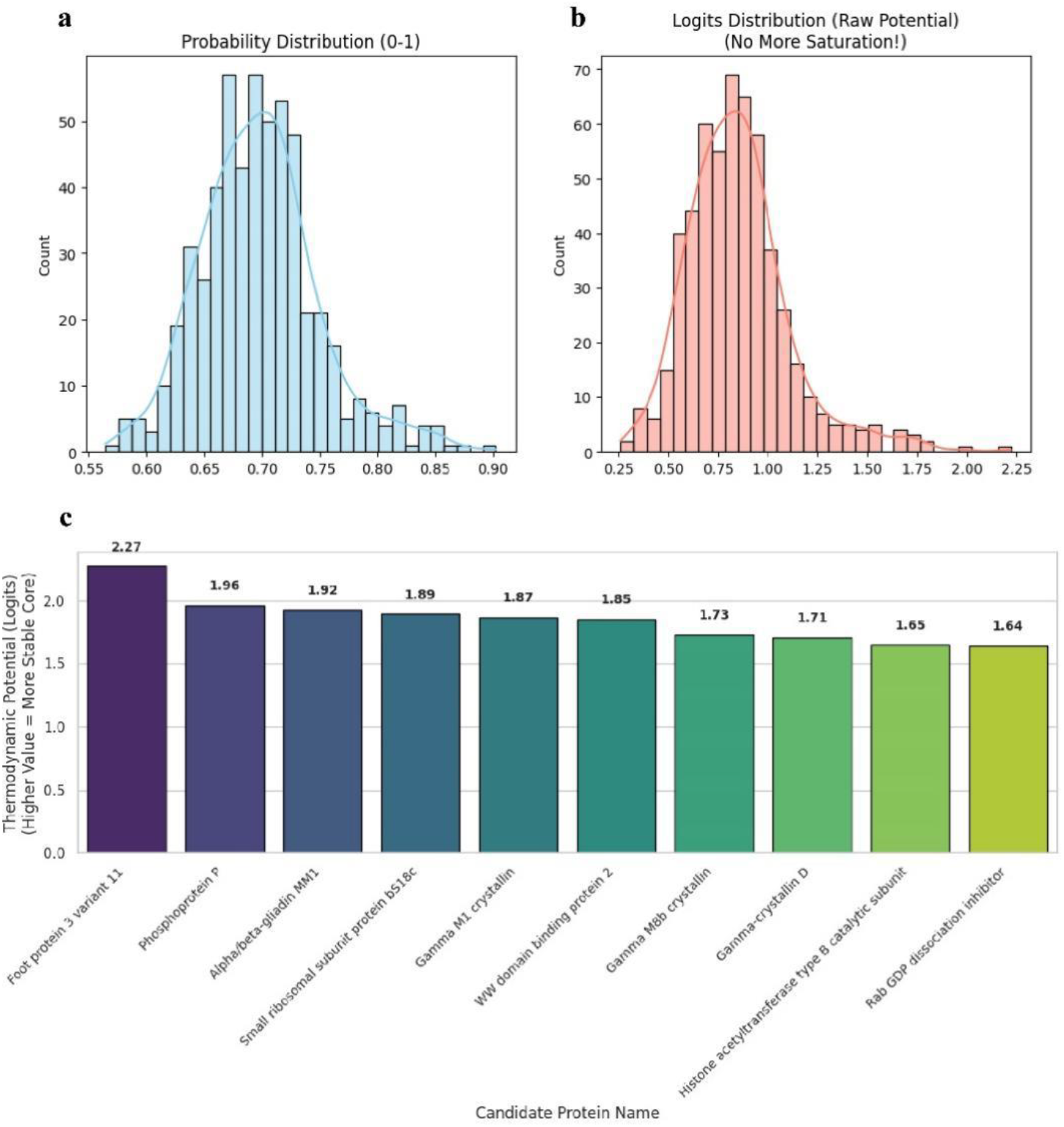
Comparison of prediction score distributions for candidate membraneless organelle (MLO) proteins. **Top:** The probability distribution after processing by the Sigmoid activation function shows a significant saturation effect, with the vast majority of candidate scores clustered in the 0.95∼1.0 interval. **Bottom:** The distribution of the original output Logits (thermodynamic potential) shows obvious heterogeneity. Higher Logits values represent stronger thermodynamic stability of the phase separation core, providing higher resolution for distinguishing high-confidence candidate proteins. **Fig. 5c English names of candidate membraneless organelle proteins** Foot protein 3 variant 11, Phosphoprotein P, Alpha/beta-gliadin MM1, Small ribosomal subunit protein bS18c, Gamma M1 crystallin, WW domain binding protein 2, Gamma M8b crystallin, Gamma-crystallin D, Histone acetyltransferase type B catalytic subunit, Rab GDP dissociation inhibitor

## 3 Discussion

This study, through three stages of model iteration, successfully progressed a basic sequence classifier into an artificial intelligence system capable of discovering physical laws and predicting novel membraneless organelles (MLOs). This progression itself is a vivid illustration of “how artificial intelligence reshapes the research paradigm of phase separation.”

First, the correlation shown by the first-stage model with the content of the key phase-separating proteins phenylalanine/tyrosine (F/Y) is not merely a performance metric but direct evidence that the AI model can capture and quantify key protein-protein interactions. This result is consistent with the conclusion in existing literature that π-πinteractions are the core driving force of LLPS [6], indicating the model is not a “black box.” Second, the successful correction of “model hallucination” in the second stage highlights the value of introducing adversarial thinking in bioinformatics. When training data distribution is imbalanced (e.g., dominated by highly disordered sequences), deep learning models tend to learn superficial heuristic rules rather than essential laws. By constructing “trap sequences” and conducting adversarial training, we encouraged the model to abandon the simple association of “disorder equals phase separation” and instead explore deeper “sequence grammar.” This process significantly enhanced the physical realism of the model, making its predictions closer to the essential judgments of cell biology. The most important progress is reflected in the “physics engine” constructed in the third stage [7]. By introducing UMAP (Uniform Manifold Approximation and Projection) manifold learning, we mapped the model’s hidden layer features into an interpretable “thermodynamic fingerprint space.” In this space, proteins spontaneously cluster based on the intrinsic physical similarity of their phase separation behavior (not just sequence similarity), providing a novel visualization and analysis pathway for discovering functionally related MLO protein clusters. Furthermore, by introducing non-equilibrium thermodynamic constraints and outputting thermodynamic potential (thermo_potential_logits), the model achieved a key leap from “classification” to “physical modeling.” The finally screened Top 10 candidate proteins are valuable not only for their high prediction probability but also for the thermodynamic stability they exhibit, thereby providing clearer targets for subsequent wet-lab experimental validation. Certainly, this study still has some limitations. The current model’s predictive capability is limited by the coverage of existing databases and may be insufficient in identifying extremely rare or novel phase separation patterns.

Looking to the future, the “model iteration - physical validation - new discovery” research paradigm constructed in this study can embed physical laws more deeply into the architecture of artificial intelligence models, promoting the development of research systems that can not only predict the relationship between phase separation propensity and conditions (such as pH, salt concentration, etc.) but also extend to other complex cell biophysical processes, such as protein dynamic aggregation and dynamic changes in intracellular ATP concentration. We believe that through the deep integration of artificial intelligence and cell biology, it will be possible to integrate multi-dimensional information such as genomics, transcriptomics, proteomics, epigenomics, and metabolomics, thereby more precisely analyzing cell functions

